# Marine Bacteria Associated with the Green Seaweed *Ulva* sp. for the Production of Polyhydroxyalkanoates

**DOI:** 10.1101/2020.12.05.413369

**Authors:** Rima Gnaim, Mark Polikovsky, Razan Unis, Julia Sheviryov, Michael Gozin, Alexander Golberg

## Abstract

The biosynthesis of polyhydroxyalkanoate (PHA) biopolymers from certain marine microbes, associated with green macroalgae *Ulva* sp., has attracted significant attention. The *Ulva* sp. is abundant biomass in numerous locations around the world and could be easily cultivated by marine farming. The variety of sugars found in *Ulva* sp. homogenate could be used as a carbon source for microbial growth and PHA production. In this work, we isolated and explored a series of bacterial strains that function as potential producers of P(3HB), utilizing a range of common sugars found in *Ulva* sp. Analysis of 16S rDNA gene-sequence revealed that the PHA-producing bacteria were phylogenetically related to species of the genus *Cobetia, Bacillus, Pseudoaltermonas*, and *Sulfitobacter*. The highest-yield of P(3HB) was observed in the case of new *Cobetia* strain, *C. amphilecti*, with up to 61% (w/w) in the presence of mannitol and 12% (w/w) on *Ulva* sp. acid hydrolysate as a substrate.

## Introduction

Limited petroleum resources and their significant environmental impacts have led to an increase in biopolymers development from renewable resources (Li et al., 2016). Among these polymers are polyhydroxyalkanoats.

Polyhydroxyalkanoates (PHAs) are prospective substitutes for petrochemical-derived polymers due to their biodegradability, sustainability, and versatile thermal and mechanical properties (Grigore et al., 2019; Muhammadi et al., 2015). PHAs are intracellular microbial aliphatic polyesters, which are synthesized by numerous organisms as carbon and energy storage in intercellular granules (Grigore et al., 2019). The PHAs are usually produced as a response to environmental stresses such as nutrient limitation (Kasan et al., 2015).

To date, around 150 various chemical structures of PHA were reported (Sagong et al., 2018). Poly-β-hydroxybutyrate (P(3HB)) gained more recognition due to its unique physio-mechanical properties. Thus, it offers great potential for use in various industrial applications in agriculture, food packaging and bio-medical fields (Mostafa et al., 2020a). The development of desirable PHA polymers from a widespread microbial resource for industrial purposes are being investigated (Kourmentza et al., 2017). Recently, marine microbial strains such as *Alteromonas, Bacillus, Pseudomonas* spp., *Cupriavidus* spp. (Możejko-Ciesielska and Kiewisz, 2016) have gained a lot of attention, they can produce superior PHA polymers because of the stressed marine conditions they live in (Mostafa et al., 2020b).

Although many bacterial species have been identified to produce PHA, the potential to discover and identify novel marine species isolated from green macroalgae with vastly superior production capacity remains untapped. Besides, optimization of bacteria growth and PHA accumulation using various carbon sources presents an essential component for the commercialization of these biopolymers (Sangkharak and Prasertsan, 2012).

Marine macroalgae or seaweeds, especially *Ulva* sp. are one of the most attractive biomass for exploring PHA production by their associated bacteria due to macroalgae abundance in many ecosystems on earth (Wei et al., 2013). This type of seaweeds offers a lot of environmental and biotechnological benefits comparing to terrestrial crops. For example, they are easily accumulated in many areas around the world; they don’t require harsh agronomical treatments, they have high growth rates and high polysaccharide content (Robic et al., 2009) making them a stellar for large-scale production (Gajaria et al., 2017; Jones and Mayfield, 2012).

Numerous studies have described the biosynthesis of a wide range of valuable materials such as biogas, butanol, and ethanol by fermentation of seaweed (Ashokkumar et al., 2017; Leaves and Based, 2019; Scientific et al., 1983; Wise et al., 1979). However, recently seaweed has been explored as a potential substrate for PHA production. Studies have shown that bacteria accumulated PHA in a medium containing brown algae (Azizi et al., 2017; Moriya et al., 2020; Muhammad et al., 2020), red algae (Alkotaini et al., 2016; Bera et al., 2015; Sawant et al., 2018), and green seaweed *Ulva* sp. (Ghosh et al., 2019). Our research group has demonstrated that the *Ulva* sp. hydrolysate is a promising feedstock for PHA production using Haloferax *mediterranei* (Ghosh et al., 2019).

In the present study, more than one hundred strains of bacteria isolated from green macroalgae *Ulva* sp. were evaluated for their capability to manufacture PHAs with various supplemented fermentative substrates found to be in macroalgae, e.g., glucose, fructose, galactose, mannitol, mannose, arabinose, rhamnose, glucuronic acid, and xylose. A total of thirty-one bacteria found to produce PHA. Ten strains related to genus *Cobetia, Bacillus, Pseudoaltermonas*, and *Sulfito-bacter*, which showed high PHA yields among the isolates, were further investigated. The effect of the type of supplemented sugars on the growth and PHA productivity of the strains was studied. Furthermore, the effect of bacteria co-culture and mixed substrates on the production of PHA was investigated. Also, 16S rRNA sequence identification of several isolated bacteria was performed. Finally, the ability of the strain *Cobetia* 105 to produce PHA on *Ulva* sp. acid hydrolysate, was demonstrated. This study could contribute to the understanding of the diversity of bacteria, associated with marine macroalgae, in terms of PHA productivity and bacteria strains.

## Materials and Methods

### Chemicals, instruments and media

For bacterial cultivation on plates, Agar powder (2% w/v) (Difco, USA) was dissolved in a medium with Marine Broth (Beit Dekel, Israel) containing (per L) 19.4 g NaCl, 3.24 g MgSO_4_·7H_2_O, 5.0 g Peptone, 8.8 g MgCl_2_·6H_2_O, 1.8 g CaCl_2_, 1 g yeast extract, 0.55 g KCl and 0.16 g NaHCO_3_ (pH 7.6). The supplemented sugars (glucose, fructose, galactose, mannitol, mannose, arabinose, rhamnose, glucuronic acid, and xylose) were purchased from Sigma-Aldrich (Israel). Nile Blue (Sigma-Aldrich, Israel) for staining of PHA was used for the screening of isolated bacteria. The sugar’s solutions were filtered through a 0.22 µm pore membrane microfilter (CSI, Israel). Bateria in liquid cultures was grown in aerobic flask bottles (175 mL) in a shaking incubator.

### Growth of the green macroalga *Ulva* sp

The growth of *Ulva* sp. was carried out by adding 20 gram of fresh *Ulva* sp. in 40 mL cylindrical, sleeve-like seaweed photobioreactor (MPBR, Polytiv, Israel) (Chemodanov et al., 2017) in a seawater medium containing 3.7% w/v of dried Red Sea salt (Red Sea Inc, IS), ammonium nitrate (NH_4_NO_3_, Haifa Chemicals Ltd, Israel) and phosphoric acid (H_3_PO_4_, Haifa Chemicals Ltd, Israel). The final concentration of nitrogen (N_2_) and phosphorus (P) in the medium were 6.4 g m^−3^ and 0.97 g m^−3^, respectively. The pH, temperature and flow rate were controlled as stated in our earlier work (Chemodanov et al., 2017).

### Acid hydrolysis of the green macroalga *Ulva* sp

*Ulva sp*. was dried at a temperature of 40°C after harvesting. Subsequently, the dried biomass was crushed with an electric grinder (Grinding machine, Henan Gelgoog Machinery GG9FZ-19) to obtain fine powdered *Ulva* sp. The acid hydrolysis was performed as described in our previous study (Id et al., 2020). Briefly, 45 grams of dry powdered *Ulva* sp. were added to 500 mL of sulfuric acid solution (2% v/v). The sample was autoclaved at 121 °C for 30 minutes. The solution was cooled, and the pH was adjusted to 6.7 by adding 117 mL of 3M NaOH solution and 80.6 mL of PBS buffer (Phosphate Buffer Saline). Subsequently, 12.2 mL of Marine Broth was added to the medium to supplement minerals and nitrogen sources, and the solution was filtered with 0.22 µm syringe-filter (Millipore, USA).

### Analysis of *Ulva* sp. acid hydrolysate by Ion chromatography

The chemical composition of the *Ulva* sp. acid hydrolysate products was determined using high-pressure ion chromatography (HPIC) via Dionex ICS-5000 (Dionex, Thermo Fischer Scientific, MA, USA). The acid hydrolysate solution was diluted in ultrapure water to reach a ratio of 1:2. The sample was then filtered with a 0.22 µm syringe filter (Millipore, USA) and added to HPIC vials (Thermo Fischer Scientific, USA). The phase flow rate was 0.25 mL/min, and the column temperature was set to 30°C. The standards used as a reference to identify and quantify the resulted monosaccharides were fructose, xylose, glucose, galactose, rhamnose and Glucuronic acid (Sigma-Aldrich, Saint-Louis, USA).

### Isolation of bacterial strains

*Ulva sp*., a green seaweed collected from the Mediterranean Sea, was used as a source of PHA-producing bacteria. The isolation of bacteria was carried out by three to six isolation rounds, up to achieving a homogenous single colony, which was detected by binocular (Figure 1). The first isolation round was done by smearing live algae thalli after it was harvested in the Israeli eastern part of the Mediterranean Sea or by smearing seawater. The first bacterial isolation round was done on plates with five different carbon sources with three different concentrations of the agar 0.7, 1 and 1.5% agar. The five media contents were: (1) natural Mediterranean seawater (SW); (2) live *Ulva* sp. (5 g wet weight) with double-distilled water (DDW); (3) *Ulva* sp. dried at 40°C and was ground with mortar and pestle, for the medium preparation was taken 1.5% of *Ulva* sp. dry weight (DW) with DDW; (4) marine broth (MB) (Marine Broth 2216, BD Difco), 3.7 g L^-1^ in DDW; (5) DDW without any carbon source. All media were autoclaved and poured into Petri dishes. The subsequent isolation rounds were done by streak-plating bacteria cultures up to isolate a single colony. All isolation rounds after the first round were done on MB plates (1.5% agar). Finally, 110 isolated bacteria colonies were transferred to 2 mL liquid marine broth (3.7 g·L^-1^) and kept for overnight at 32°C in a shaker incubator (180 RPM, Incu-Shaker Mini, Benchmark Scientific). The bacteria were stored in glycerol (final concentration of 25%) at -80°C.

**Figure 1.**
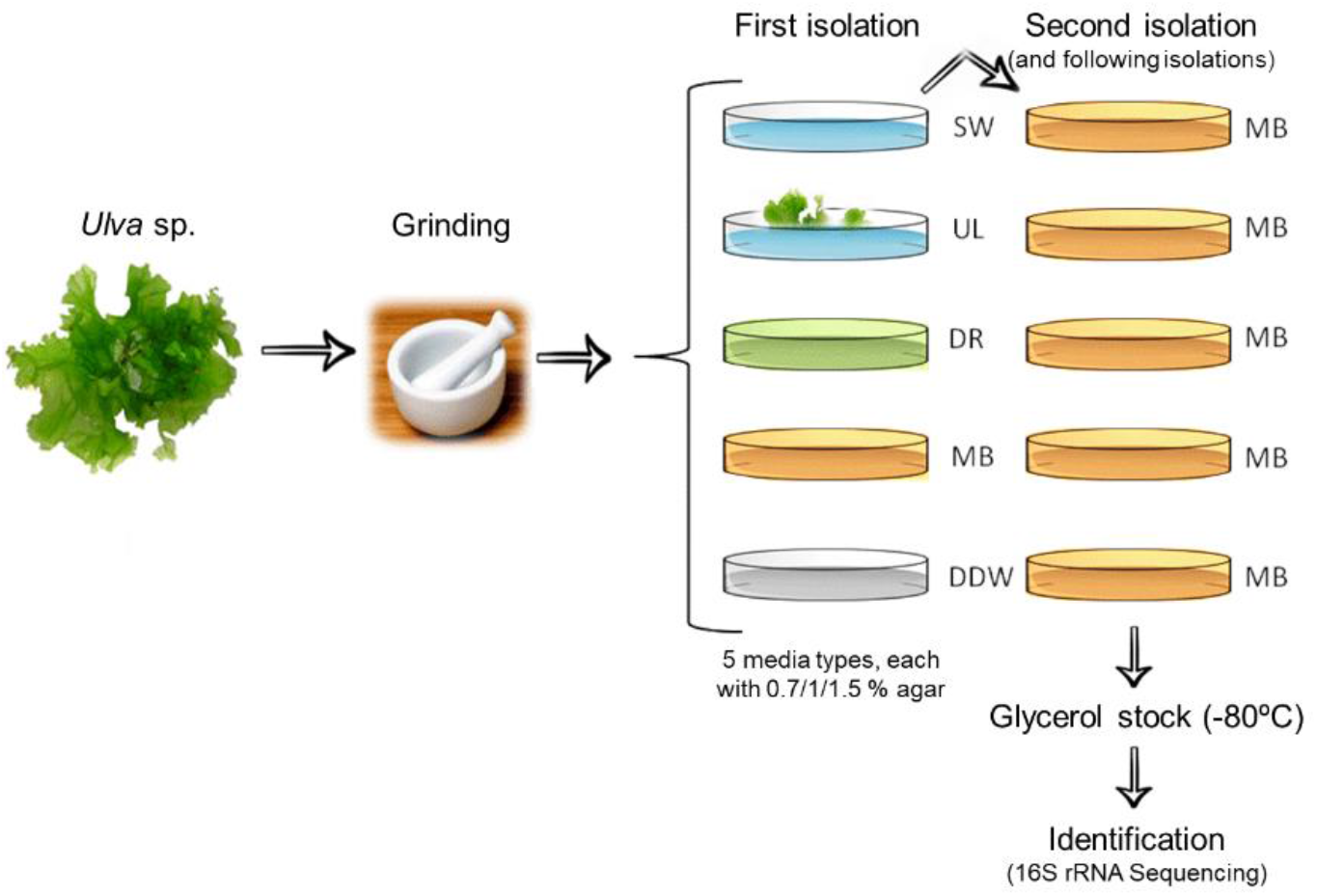
Illustration of the bacterial isolation from *Ulva* sp. The *Ulva* associated bacteria were isolated after *Ulva* sp. was ground and streaked on different plates. The isolated bacteria were stored and genetically identified.

### Screening of bacteria utilizing different sugars for PHA production

All of 110 bacterial isolates were tested for PHA production using Nile Blue A staining method. The bacteria isolates were cultivated on agar plates containing MB and the selected sugar (2% w/v) and incubated for four days at 32°C. Nile Blue A (0.5 *μ*g/mL) was directly added to a rich Marine Broth agar medium; thus, the bacterial cells were grown in the presence of the dye. Subsequently, the bacteria were exposed to UV illumination (320 nm) using the ENDURO™ GDS Gel Documentation System (Labnet International, Inc. Israel). This technique allowed rapid screening of the viable colonies for PHA production and considered to be a powerful tool for distinguishing between PHA-negative and PHA-positive strains. The bacteria that have shown a bright white fluorescence on irradiation with UV light were selected as potential PHA accumulators. The selected bacteria were repeatedly grown on different sugars in Marine Broth plates, and the accumulation of PHA on each sugar was also examined by Nile Blue staining. All experiments were carried out in triplicates.

### Molecular identification of the isolates

PHA-positive bacteria were genomically identified to the genus level using 16S sequencing profiling. For strain identification, genomic DNA extraction was performed, a colony of each bacterial strain was transferred into a 2 mL sterile tube containing distilled water. The samples were then centrifuged for 3 minutes at 10000 RPM and heated for 10 minutes at 100°C in an Eppendorf Thermomixer C (Thermo Fisher Scientific, USA) to lyse the bacterial cells. The supernatant of the sample, which contains the DNA fragments, was obtained, and the cell pellet was discarded. The microbial DNA was purified using the Exo-sap DNA Clean-Up Kit (Sigma-Aldrich, Israel) using 5 L aliquot of the supernatant. The 16S rDNA was amplified by PCR using standard protocols (Wang et al., 2011) based on the primers data shown in Table 1. The PCR product was purified by Exo-sap clean up kit. Sequencing of 16S rDNA was carried out by TAU genomic unit, and a homology search of the databases was performed using the BLAST. A phylogenetic tree was constructed using the neighbor-joining DNA distance algorithm (Saitou and Nei, 1987) using Mega 5. The resultant tree topologies were evaluated by bootstrap analysis of neighbor-joining data sets based on 100 resamplings.

**Table 1.**
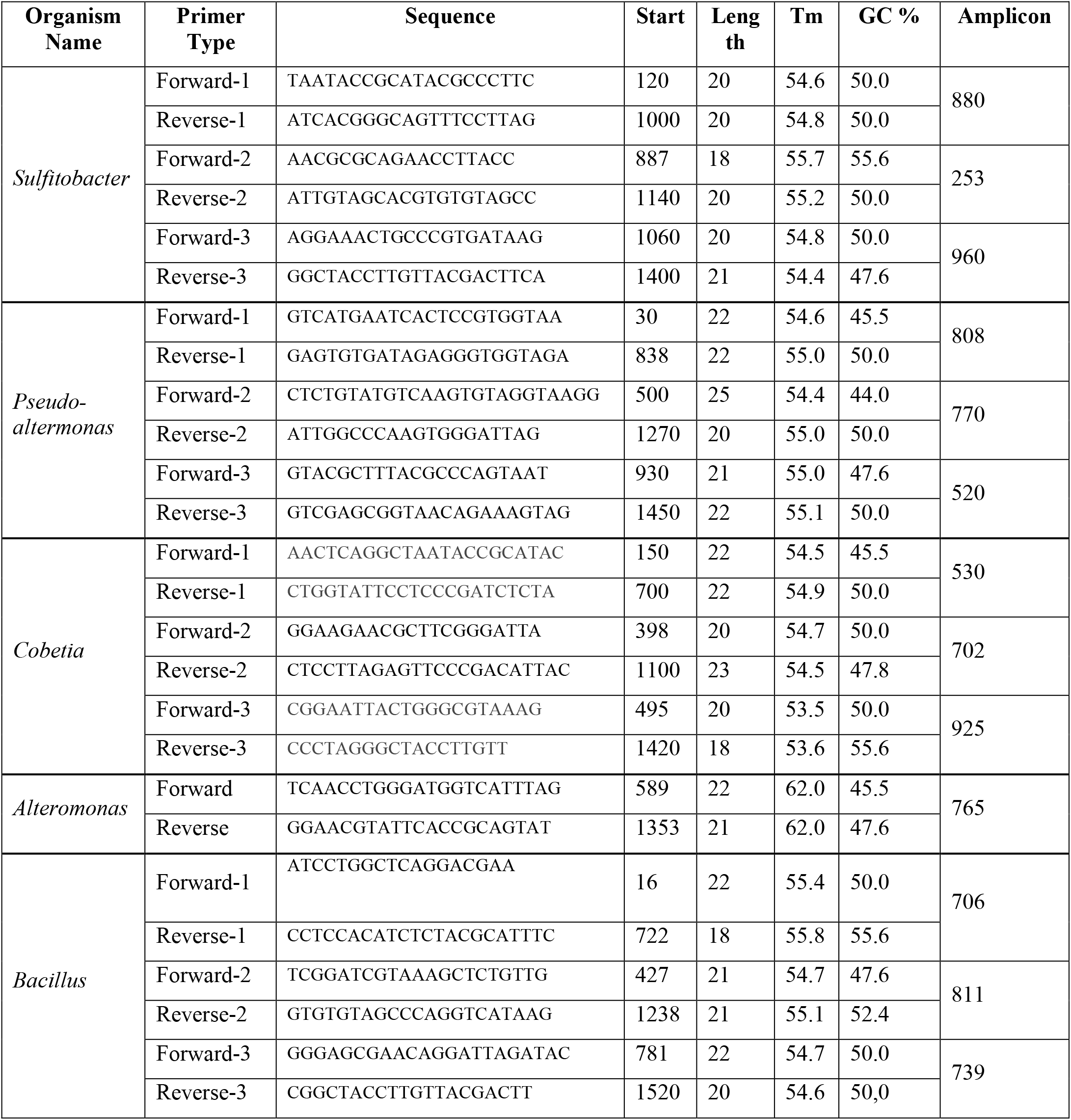
PCR primers and conditions used for bacteria identification

### Cultivating of PHA-producing isolates in liquid media with different sugars

Starters of the selected PHA-positive bacteria were prepared by adding one bacterial colony into Marine Broth (MB) medium and were incubated for 18 hours at 37°C. The bacteria starters were poured into a single sterile bottle. 1.75% of MB media (900 mL) was prepared and autoclaved. The selected carbon source (2% w/v) was dissolved in the medium and adjusted to afford pH 7. For each treatment, a sterile glass bottle containing 135 mL of MB media was prepared. Subsequently, 15 mL of bacteria were added from the bacteria inoculum to the medium (total volume of 150 mL). The content of the bottles was then appropriately mixed, and the 150 mL solutions were divided to afford three portions of 50 mL solutions. The cultures were grown under aerobic conditions in a shaking incubator at 32°C with a rotational speed of 90 rpm for four days. The bacterial growth was examined by measuring OD 600. The resulting biomass was collected by centrifugation at 4500 g for 30 min in a swing rotor centrifuge (Rotanta 420R, Hettich Instruments LP, USA), rinsed twice with a saline solution followed by 15 min centrifugation, dried in an oven at 45°C for 24h until a constant weight was obtained. The DW biomass and %DW per fermentation volume was calculated. PHAs were extracted and analyzed by GC-MS and ^1^H-NMR. All experiments were carried out in triplicates.

### PHA production by bacterial combinations on sugar mixtures

The best PHA-producing bacteria were chosen to study the effect of bacteria combination and sugar mixture on PHA production. Starters of the selected PHA-positive bacteria were prepared by adding one bacterial colony into Marine Broth (MB) medium and were incubated for 18 hours at 37°C following the previously mentioned procedure. The bacteria starters were poured equally (5 mL each bacteria) into a sterile bottle. MB media was prepared and autoclaved. The selected carbon source was added to the media (2% w/v for each sugar type) and adjusted to afford pH 7. For each treatment, a sterile glass bottle containing 135 mL of MB media was prepared. Subsequently, 15 mL of bacteria were added from the bacteria inoculum to the media to yield 150 mL of solution. The bottles were mixed properly, and the 150 mL solutions were divided equally into three 50 mL solutions. The cultures were grown under aerobic conditions in a shaker (90 rpm) at 32°C for 4 days. The bacterial growth was examined by measuring OD 600. The resulting biomass was collected by centrifugation at 4500 g for 30 min in a swing rotor centrifuge (Rotanta 420R, Hettich Instruments LP, USA), rinsed twice with a saline solution followed by 15 min centrifugation, dried in an oven at 45°C for 24h until a constant weight was obtained. The DW biomass and %DW per fermentation volume was calculated. PHAs were extracted and analyzed using GC-MS and ^1^H-NMR. All experiments were carried out in triplicates.

### PHA production by *Cobetia* 105 on *Ulva* sp. acid hydrolysate

Starters of Cobetia isolate no. 105 were prepared by adding one bacterial colony into Marine Broth (MB) medium and were incubated for 18 hours at 37°C following the procedure mentioned above. The bacteria starters were poured into a single sterile bottle. The selected carbon source was added to the *Ulva* sp. hydrolysate media (2% w/v). A sterile glass bottle containing 135 mL of hydrolysate media was prepared. Subsequently, 15 mL of bacteria were added from the bacteria inoculum to the media to yield 150 mL of solution. The bottles were then mixed properly, and the 150 mL solutions were divided equally into three 50 mL solutions. The cultures were grown under aerobic conditions in a shaker (90 rpm) at 32°C for 4 days. The bacterial growth was examined by measuring OD 600. The resulting biomass was collected by centrifugation at 4500 g for 30 min in a swing rotor centrifuge (Rotanta 420R, Hettich Instruments LP, USA), rinsed twice with a saline solution followed by 15 min centrifugation, dried in an oven at 45°C for 24h until a constant weight was obtained. The DW biomass and %DW per fermentation volume was calculated. PHAs were extracted and analyzed using GC-MS and ^1^H-NMR. All experiments were carried out in triplicates.

### Characterization and quantification of PHA by GC-MS

PHAs were analyzed after direct acid-catalyzed trans-esterification with methanol of the dried bacteria (DB). The tested samples of DB (10-30 mg) were added to a mixture of chloroform (1.0 mL), benzoic acid (1.0 mg, an internal standard, BA), methanol (2.0 mL) and concentrated H_2_SO_4_ (0.5 mL). The suspension was heated at 90°C with magnetic stirring for overnight in a closed vial. The reaction mixture was cooled to room temperature and treated with a cooled saturated NaCl solution (15 mL) and chloroform (10 mL). Anisole (1.0 mg, an external standard, AN) and 2,4-dimethylanisole (1.0 mg, an external standard, DMA) were added to the mixture. The organic phase was washed twice with water, separated, dried over anhydrous sodium sulfate and concentrated under vacuum to obtain 1 mL solution. GC-MS was used to analyze the PHA methanolysis products and their chemical composition. GC-MS analysis was performed using a Thermo Trace 1310 GC, equipped with a TG-SQC GC capillary column (15 m, 0.25 mm i.d., 0.25 µm film thickness) and a mass spectrometer ISQ LT as the detector. The carrier gas was helium at a flow rate of 1.2 mL/min. The column temperature was initially 50°C for 1 min, then gradually increased to 200°C at 10°C/min, and finally increased to 285°C at 20°C/min. For GC-MS detection, an electron ionization system was used with ionization energy of 70 eV. The samples were diluted 1:1000 (v/v) with ultra-pure hexane, and 1.0 µL of the diluted samples (8 ng/1 µL) was injected automatically in split mode. Injector and detector temperatures were set at 250°C. All experiments were carried out in triplicates.

### ^1^H NMR analysis

All samples were dissolved in deuterated CDCl_3_ prior analysis (5 mg/mL). Each sample was shaken vigorously till complete dissolution was achieved, and about 0.5 mL of it was transferred into an NMR tube for analysis and run ^1^H-NMR with Pulse Program zg30 on Bruker AVANCE III 500 MHz NMR Spectrometer with 5 mm PABBO-BB probe and Topspin 3.0 software.

### Statistical Analysis

The results were statistically analyzed using Excel and GraphPad prism 8 for data management and quantitative analysis. One-way and two-way ANOVA using Tukey and Holm-Sidak’s multiple comparison tests were performed for analyzing standard deviation, means and statistical significance for PHA yield and DCW concentration.

## Results

### PHA accumulation by *Ulva* sp. associated bacterial strains utilizing different sugars

A total of 110 bacteria were isolated from *Ulva sp*. and screened for PHA production by using Nile Blue A staining method. All positive-PHA strains exhibited a white fluorescent emission on agar plates containing different sugars under UV light. For example, Figure 2 shows the diversity of PHA production of *Cobetia* isolate no. 104 that produces PHA mainly in mannitol, fructose, galactose, and glucose, while no PHA was observed with *Cobetia* isolate no. 104 in the presence of other sugars. It is important to emphasize that all tested bacteria did not produce PHA when grown on MB alone as a control. Based on fluorescence staining, 28 bacteria were found to accumulate PHA to a different extent in the presence of glucose, fructose, mannitol, and galactose. The sugar substrate with the highest number of bacteria that produce PHA is glucose with 27 different strains, followed by fructose with 24 strains, then mannitol and galactose with 17 strains (Table 2).

**Table 2.** List of bacterial isolates which showed a white light fluorescence under UV light when grown on different sugars. The white fluorescence indicates the accumulating of PHA (Oshiki et al., 2011).

| Bacteria's no. | Bacteria genus | Sugar type |  |  |  |  |  |  |  |  |
| --- | --- | --- | --- | --- | --- | --- | --- | --- | --- | --- |
|  |  | Gal | Mat | Fru | Ara | Mas | Glu | Rha | GA | Xyl |
| 49 | *Sulfitobacter* | + |  | + |  |  | + |  |  | + |
| 3 | *Bacillus sp.* | + | + | + |  |  | + |  |  |  |
| 25, 26, 27 | *Uncultured Altermonas* |  | + | + |  |  | + | + |  | + |
| 28 | *Altermonas* |  | + | + |  |  | + | + |  | + |
| 52, 56 | *Unclassified vibrio* |  | + | + |  |  | + |  |  |  |
| 41 | *Vibrio sp.* |  | + | + |  |  | + |  |  |  |
| 6, 37 | *Sulfitobacter sp.* |  |  |  |  |  | + |  |  | + |
| 68, 80, 81, 85 | *Unclassified Microbacteria* | + |  | + | + | + | + |  |  | + |
| 86 | *Unclassified Microbacteria* | + |  | + | + | + | + | + |  | + |
| 75-76, 92, 104-105, 107 | *Cobetia* | + | + | + |  |  | + |  |  |  |
| 13 | *Pseudoaltermononas* | + | + | + |  | + | + |  |  |  |
| 71 | *Pseudoaltermononas* | + |  | + | + | + | + | + |  |  |
| 14 | *Pseudoaltermononas* | + | + | + |  |  | + |  |  |  |
| 65 | *Cobetia* | + | + | + |  |  | + |  |  |  |
| 82 | *Sulfitobacter sp.* |  |  |  |  |  | + |  |  |  |
| 48 | *Sulfitobacter sp.* |  |  |  |  |  |  |  |  | + |

+ Positive-PHA. Gal-galactose; Mat-mannitol; Fru-fructose; Ara-arabinose; Mas-mannose; Glu-glucose; Rha-rhamnose; GA-glucuronic Acid; Xyl-xylose.

**Figure 2.**
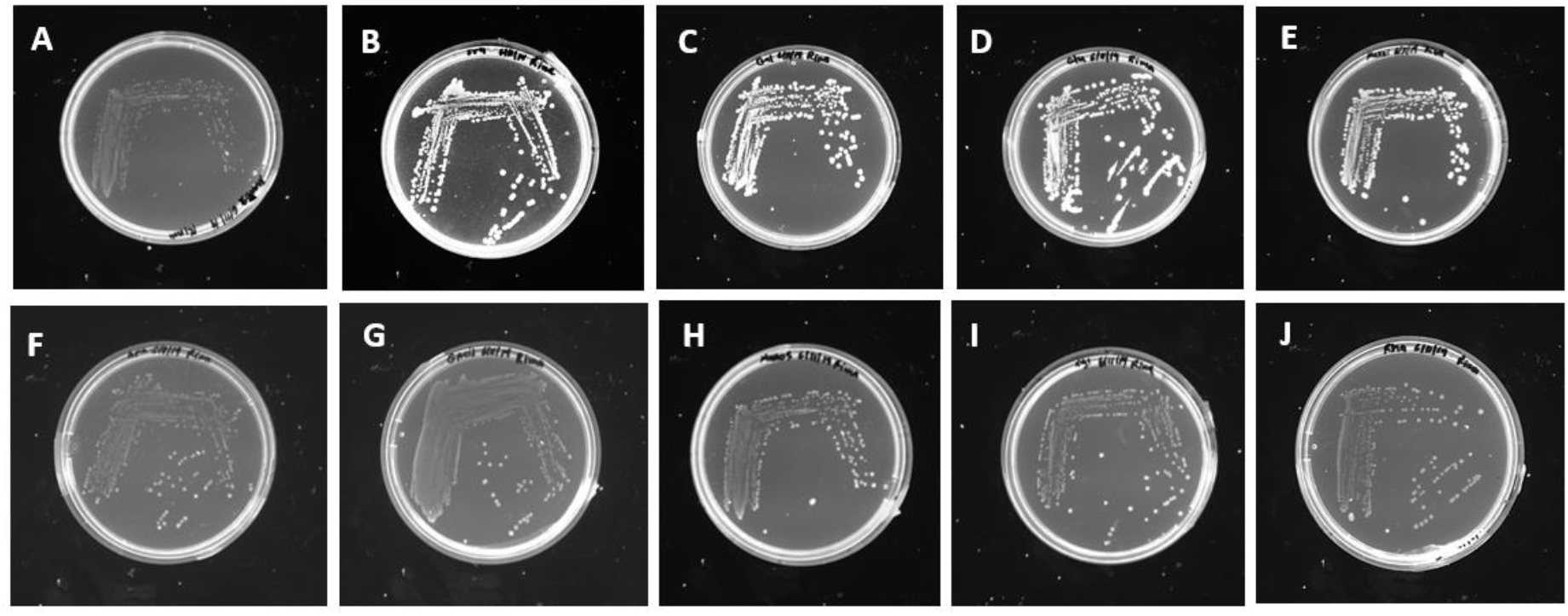
Screening of *Cobetia* (no. 104) for PHA production on different sugars under UV light: **A**-negative control, **B**-fructose, **C**-galactose, **D**-glucose, **E**-mannitol, **F**-arabinose, **G**-glucuronic acid, **H**-mannose, **I**-xylose, **J**-rhamnose. Fluorescence indicates the presence of PHA accumulation inside the bacterial cells. *Cobetia* (no. 104) utilize mannitol, fructose, galactose and glucose for PHA production while no PHA is observed on other sugars.

### PHA-Producing Bacteria Identification Using 16S rRNA Gene and Phylogenetic Analysis

Molecular identification of the isolates was carried out by the sequencing of 16S rRNA gene. Amplification of bacterial genomic DNA by primers yielded 1400-1500 bp fragments (Figure 3). The bacteria were found to be within the genus of *Cobetia, Bacillus, Sulfitobacter* and *Pseudoaltermonas*. The phylogenetic relationship among the *Cobetia* isolates is provided in Figure 4. *Cobetia* no. 65, *Cobetia* no. 92, and *Cobetia* no. 104 were found to have 16S rRNA similarity and close relation to *Cobetia amphilecti. Cobetia* no. 105, *Cobetia* no. 75, *Cobetia* no. 76. On the other hand, *Cobetia* no. 107 were found to have a close genomic characterization and likely related to both *C. pacifica* and *C. litoralis*. Besides, *Cobetia* isolates found to have a strong evolutionary relationship with *Halomonas*, as was suggested by Arahal et al. (Arahal et al., 2002). Most of these bacteria are PHA producers. *Bacillus* isolate no. 3, also found to have a close genomic relationship to *B*. cereus, *B. mobilis, B. pacificus* and *B. thuringiensis* with 98% identity. Two or more distinct *Bacillus* species may possess identical 16S rDNA sequences (Ash et al., 1991; IJsselmuiden and Faden, 1992). Additional taxonomic studies on the isolates showed that isolate no. 48 has a genomic relationship to *Sulfitobacter* sp. and isolate no. 71 has a genomic relationship to *Pseudoaltermonas* sp.

**Figure 3.**
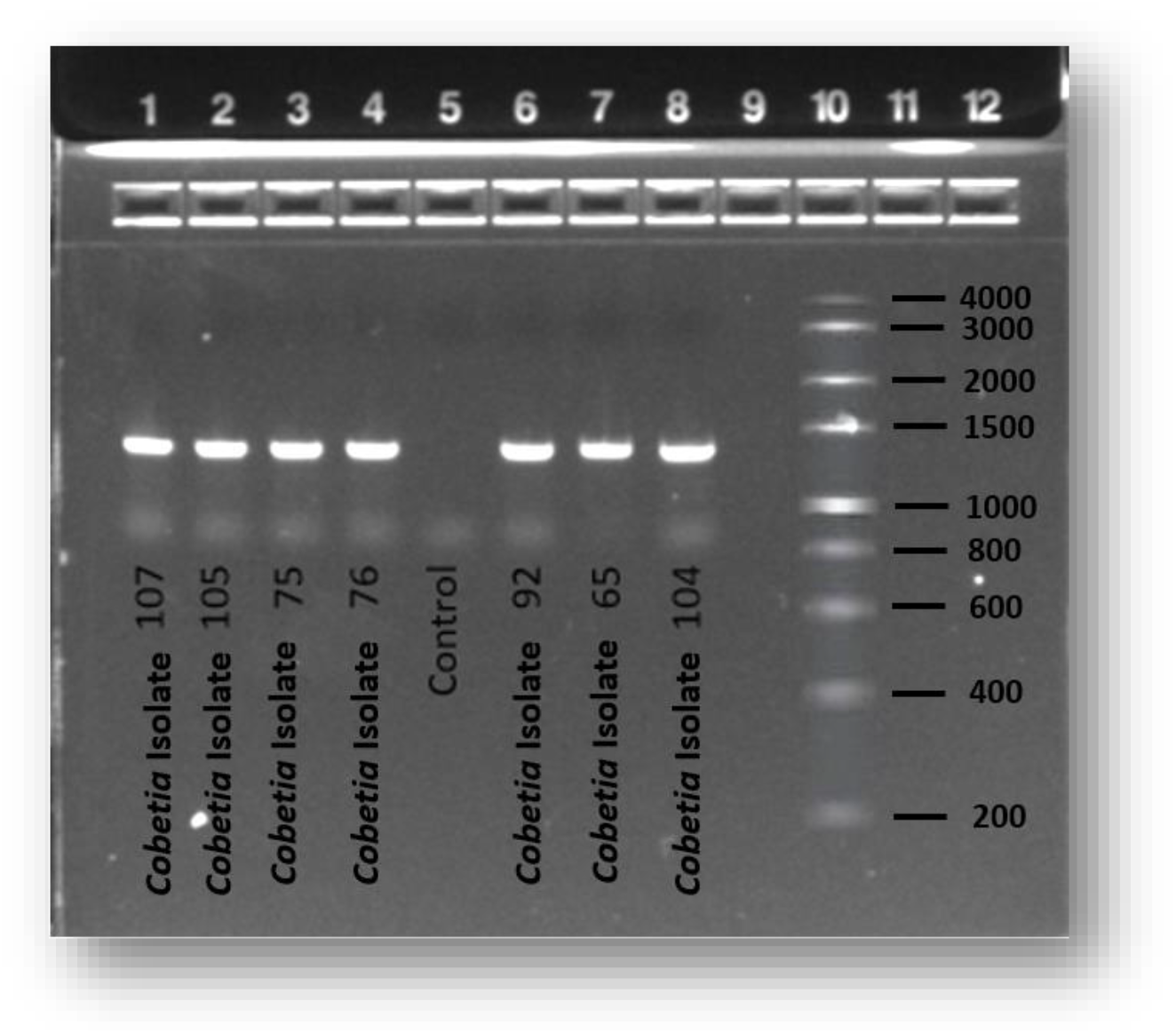
Agarose gel electrophoresis represents the amplicon of 16S rRNA gene of strains isolated from seaweeds associated bacteria.

**Figure 4.**
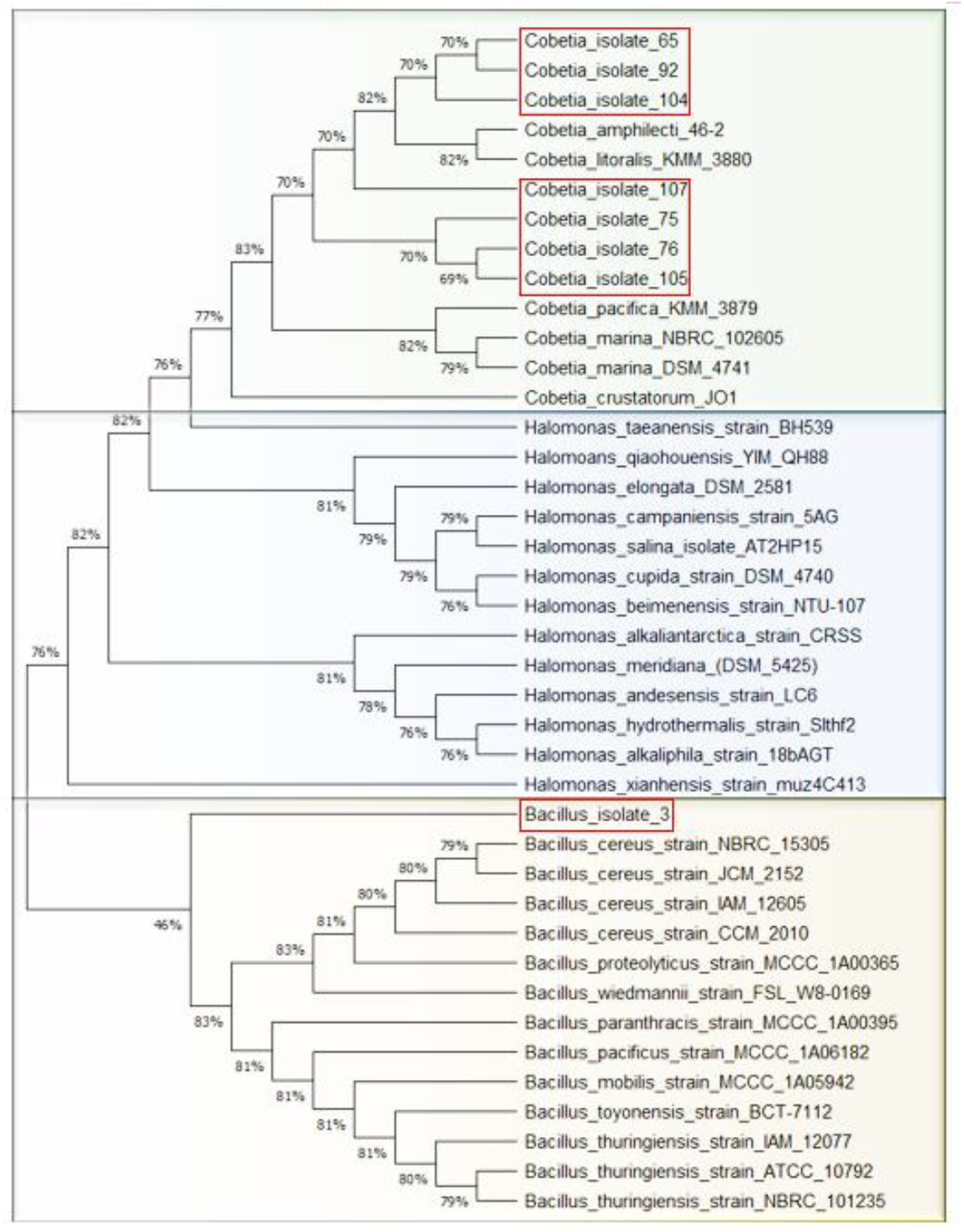
Phylogenetic relationships among the isolates drawn by MEGA 5 with 100 resamplings. The bacteria in bold were isolated in the current study.

### Chemical structure and amount of the produced P(3HB) by *Cobetia, Bacillus, Pseudoaltemonas and Sulfitobacter*

The results of the produced methylated ester derivatives obtained by acid methanolysis of PHA showed mainly two large peaks corresponding to methyl-3-hydroxybutyrate (M3HB, R_t_=3.15 min), and methyl-3-methoxy-butanoate (M3MB, R_t_=3.97 min), and a small peak corresponding to levulinic acid (LA, R_t_=5.89 min) in addition to our three standards (anisole-ANS R_t_=4.67 min; methyl benzoate-MB R_t_=7.03 min and 2,4-dimethylanisole-DMA R_t_=7.49 min) as shown in a typical GC-MS chromatogram in Figure 5. The presence of M3HB and M3MB monomers indicated that the PHA polymer is mainly P(3HB).

**Figure 5.**
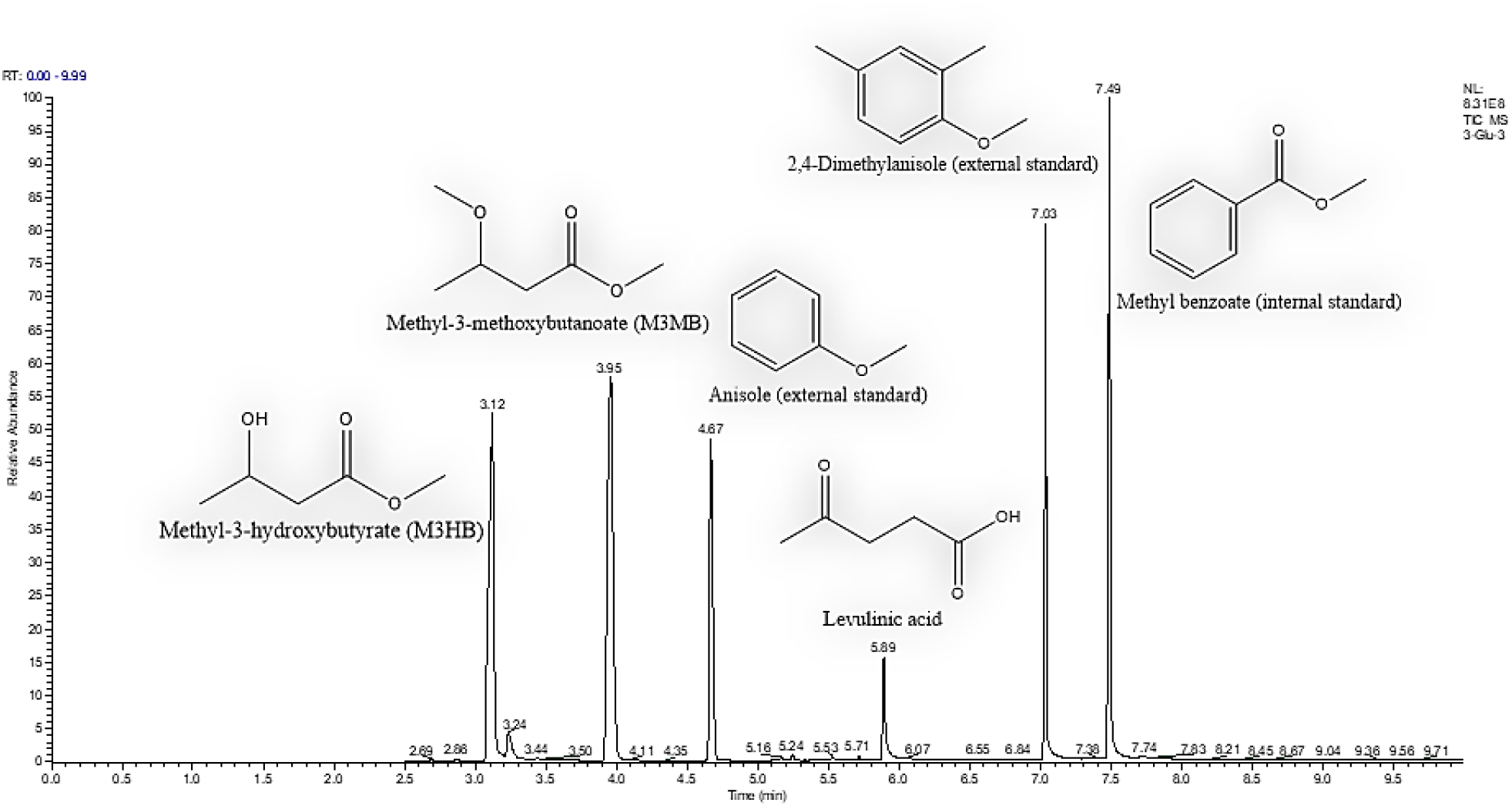
GC-MS chromatogram of methylated derivatives of 3HB (M3HB and M3MB), levulinic acid and three standards (anisole, methyl benzoate and 2,4-dimethyl anisole).

The dry cell weight (DCW, g L^-1^), P(3HB) content (%DCW), and P(3HB) yield (mg L^-1^) values obtained with different *Ulva* sp. associated bacteria grown on different supplemented sugars are presented in Table 3. Cell growth of 1.14 g L^-1^ and 1.96 g L^-1^, and P(3HB) content yield of 10.03% and 13.97% were obtained when *Bacillus* was grown in a medium containing fructose and glucose, respectively. *Sulfitobacter* produced 7.73% of P(3HB) and 2.54 g L^-1^ of DCW when it was grown in medium containing mannitol. A DCW of 6.63 g L^-1^ and 1.06 g L^-1^ and a P(3HB) production of 17.11% and 11.83% were obtained with *Cobetia* isolate no. 65 grown in medium containing mannitol and galactose, respectively. *Pseudoaltermonas* produced 7.46% of P(3HB) with DCW of 2.54 g L^-1^ when grown in medium containing fructose, while no PHA was produced on other sugars. The highest DCW of *Cobetia* isolate no. 75 was obtained when it was grown in a medium containing mannitol or glucose (4.72 g L^-1^ and 3.72 g L^-1^, respectively), and the highest P(3HB) production was achieved with mannitol (18.56%) and glucose (20.91%). *Cobetia* isolate no. 104 produced the highest amount of P(3HB) in fructose (23.39%). *Cobetia* isolate no. 105 produced 61% of P(3HB) in mannitol with 4.58 g L^-1^of DCW. *Cobetia* isolate no. 107 produced the highest P(3HB) amount when it was grown in fructose (27.45%) with a DCW of 3.53 g L^-1^. The highest P(3HB) amount using *Cobetia* isolate no. 92 was obtained in mannitol (8.91%) with 4.5 g L^-1^ of DCW.

**Table 3.** Microbial production of PHA from different supplemented sugars. A total of 10 bacteria were analyzed on different sugars for PHA production. The best bacteria with the highest PHA production were listed in the table below. DCW represents “Dry Cell Weight”. SD represents “Standard Deviation”.

| Organism Name | Sugar Type | DCW (g L<sup>-1</sup>) | PHA Yield (mg L<sup>-1</sup>) | PHA Yield (%DCW) | SD of PHA% | Monomer Composition (mol%), 3HB |
| --- | --- | --- | --- | --- | --- | --- |
| *Bacillus isolate no. 3* | Fructose | 1.14 | 114 | 10.03 | 0.76 | 100 |
|  | Glucose | 1.96 | 221 | 13.97 | 1.13 | 100 |
| *Sulfitobacter sp. Isolate no. 48* | Mannitol | 2.54 | 196 | 7.73 | 0.98 | 100 |
| *Cobetia isolate no. 65* | Mannitol | 6.36 | 125 | 17.11 | 1.41 | 100 |
|  | Galactose | 1.06 | 573 | 11.83 | 1.45 | 100 |
| *Pseudoaltermonas isolate no. 71* | Fructose | 2.54 | 189 | 7.46 | 0.64 | 100 |
| *Cobetia isolate no. 75* | Fructose | 2.08 | 125 | 8.63 | 1.15 | 100 |
|  | Galactose | 0.78 | 180 | 16.04 | 1.32 | 100 |
|  | Mannitol | 4.72 | 762 | 18.56 | 0.88 | 100 |
|  | Glucose | 3.72 | 876 | 20.91 | 0.82 | 100 |
| *Cobetia isolate no. 76* | Mannose | 3.68 | 151 | 1.89 | 0.41 | 100 |
| *Cobetia isolate no. 92* | Fructose | 3.26 | 206 | 4.45 | 0.92 | 100 |
|  | Mannitol | 4.50 | 251 | 8.91 | 0.51 | 100 |
|  | Galactose | 1.88 | 355 | 7.69 | 1.35 | 100 |
| *Cobetia isolate no. 104* | Galactose | 1.02 | 111 | 10.87 | 1.34 | 100 |
|  | Mannitol | 1.02 | 116 | 11.34 | 0.75 | 100 |
|  | Glucose | 0.82 | 131 | 16.01 | 1.12 | 100 |
|  | Fructose | 4.44 | 718 | 23.29 | 0.57 | 100 |
| *Cobetia isolate no. 105* | Mannitol | 4.58 | 574 | 61.00 | 1.23 | 100 |
| *Cobetia isolate no. 107* | Glucose | 2.03 | 231 | 11.37 | 0.95 | 100 |
|  | Fructose | 3.53 | 968 | 27.45 | 0.84 | 100 |

*Cobetia* isolate no. 75, *Cobetia* isolate no. 92, *Cobetia* isolate no. 104 and *Cobetia* isolate no. 107 produce P(3HB) mainly on galactose, mannitol, fructose, and glucose. While *Cobetia* isolate no. 65 produce P(3HB) on galactose and mannitol. *Cobetia* isolate no. 105 produce P(3HB) mainly on mannitol and fructose, and *Cobetia* isolate no. 76 produce P(3HB) mainly on fructose, glucose, and mannose.

### Effect of bacterial combination and sugar mixtures on the PHA production

Mixed culture systems were shown to produce large amounts of PHAs in a wide range of low-cost substrates (Shalin et al., 2014). We have grown the best PHA-producers on three sugar substrates with a total concentration of 2% w/v. *Cobetia* isolate no. 107, Cobetia isolate no. 104, Cobetia isolate no. 92, Cobetia isolate no. 65, Cobetia isolate no. 75 were selected to study the effect of bacteria combinations on bacteria growth and PHA production. The selected sugars were glucose, fructose and mannitol. The results in Figure 6 present the biomass amount, PHA amount and yield of all *Cobetia* strains on a mixed culture of glucose, fructose and mannitol. The highest biomass, PHA amount and PHA yield were obtained by *Cobetia* isolate no. 105 with 2.03 g·L^−1^, 712 mg·L^−1^ and 35.1% respectively. The results presented in Table 4 show that a mixed culture of different species of bacteria afforded relatively low DCW and P(3HB) yields. For example, *Cobetia* isolate no. 107 alone and *Cobetia* isolate no. 104 alone produced 27.45% w/w and 23.29% w/w of P(3HB) in fructose, respectively. However, a mixed culture of these two bacteria in fructose afforded only 10.05% of P(3HB). A similar result was observed when a bacterial combination of *Cobetia* isolate no. 65, isolate no. 75, and isolate no. 105 in mannitol was used. A yield of 11.61% of P(3HB) was obtained for the mixed bacteria compared to 17.11% for *Cobetia* isolate no. 65, 18.56% for *Cobetia* isolate no. 75, and 61% for *Cobetia* isolate no. 105. Notably, additional valuable fine chemicals were also exhibited in a very low amount, such as hexane-2,5-dione and levulinic acid.

**Table 4.**
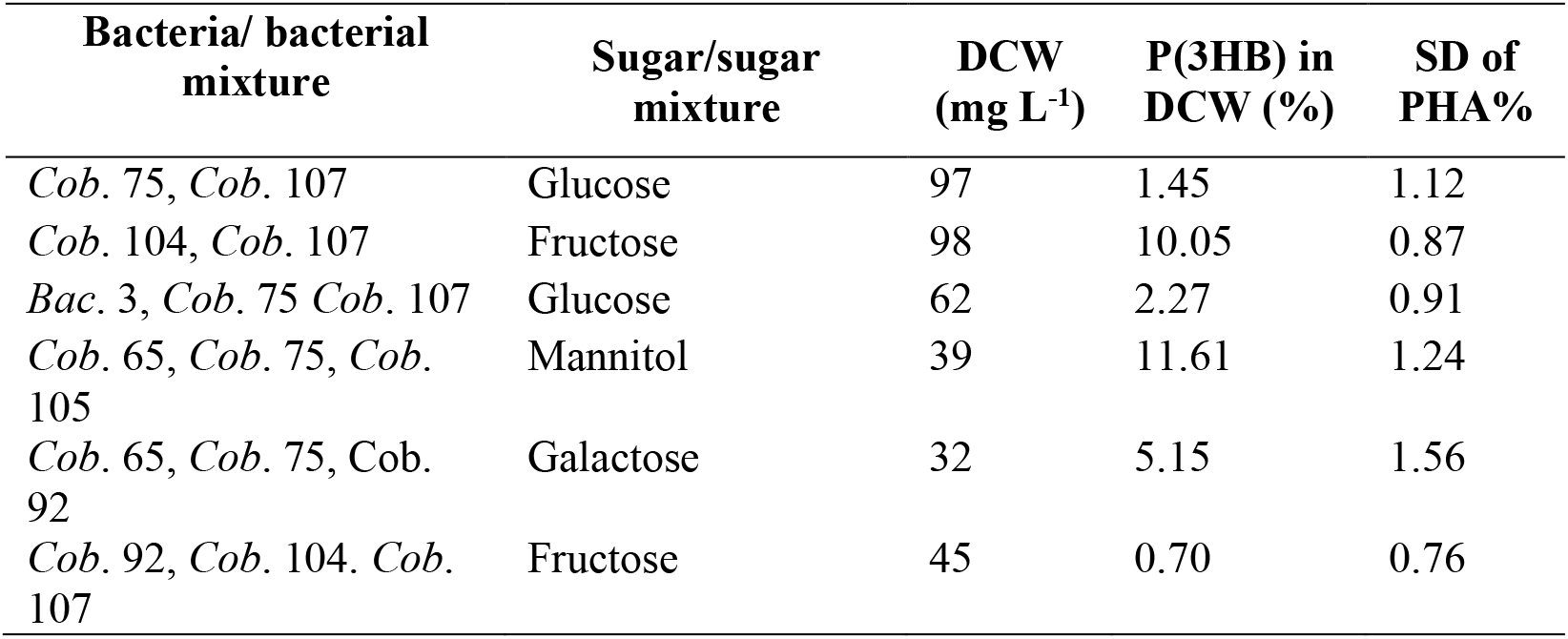
Microbial production of P(3HB) using mixed culture and mixed sugars. A Comparison between pure and mixed bacteria cultures from *Cobetia* and *Bacillus* genus are grown on single and mixture sugars substrates. DCW represents “dry cell weight”, and P(3HB) presents poly-3-hydroxybutyrate. *Cobetia* presented as Cob. and *Bacillus* presents as Bac.

| Bacteria/ bacterial mixture | Sugar/sugar mixture | DCW (mg L<sup>-1</sup>) | P(3HB) in DCW (%) | SD of PHA% |
| --- | --- | --- | --- | --- |
| *Cob.* 75, *Cob.* 107 | Glucose | 97 | 1.45 | 1.12 |
| *Cob.* 104, *Cob.* 107 | Fructose | 98 | 10.05 | 0.87 |
| *Bac.* 3, *Cob.* 75 *Cob.* 107 | Glucose | 62 | 2.27 | 0.91 |
| *Cob.* 65, *Cob.* 75, *Cob.* 105 | Mannitol | 39 | 11.61 | 1.24 |
| *Cob.* 65, *Cob.* 75, *Cob.* 92 | Galactose | 32 | 5.15 | 1.56 |
| *Cob.* 92, *Cob.* 104. *Cob.* 107 | Fructose | 45 | 0.70 | 0.76 |

**Figure 6.**
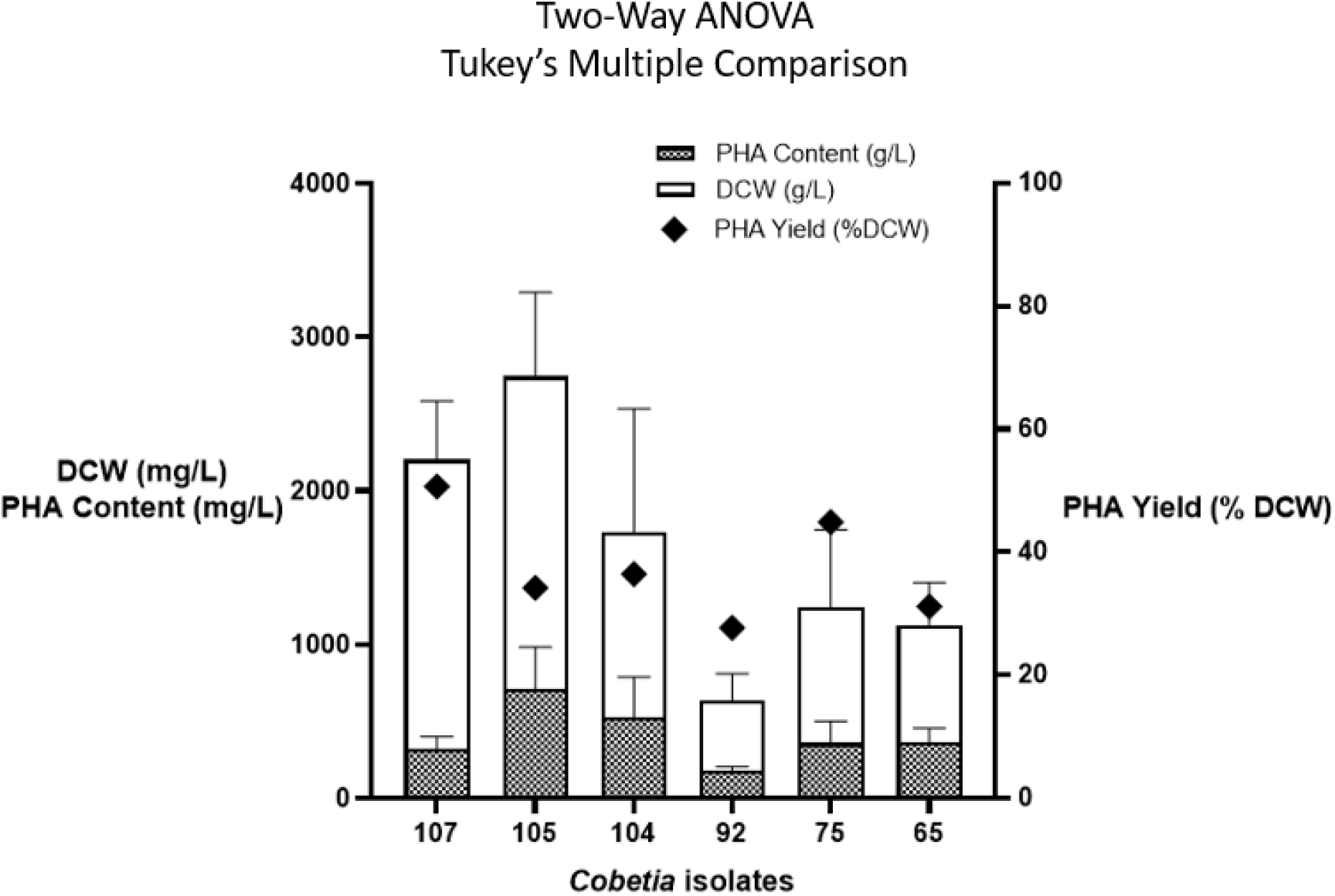
P(3HB) content, P(3HB) yield and DCW of six *Cobetia* strains (*Cobetia* isolate no. 107, *Cobetia* isolate no. 105, *Cobetia* isolate no. 104, *Cobetia* isolate no. 92, *Cobetia* isolate no. 75, and *Cobetia* isolate no. 65 grown on a mixture carbon source, i.e. glucose, fructose and mannitol. Five replicates were obtained. Two-Way ANOVA, Tukey’s multiple comparison test was performed.

### PHA production by *Cobetia* isolate no. 105 on *Ulva* sp. hydrolysate

A mixture of monosaccharides was obtained by acid hydrolysis of *Ulva* sp. which were quantified using HPIC (Table 5). The hydrolysate composed of glucose (16.1±0.8 mg/g DW), rhamnose (6.2±0.45 mg/g DW), fructose (2.8±0.41 mg/g DW), xylose (1.6±0.22 mg/g DW), galactose (1.0±0.11 mg/g DW) and glucuronic acid (1.3±0.11 mg/g DW). PHA production by *Cobetia* no. 105 on *Ulva* sp. hydrolysate was investigated (Figure 7). The results showed a biomass concentration of 1.4 ± 0.12 g·L^−1^ and PHA yield (% DCW) of 12% (w/w). The ^1^H-NMR spectrums of the PHA extracted from *Cobetia* isolate no. 105 grown on *Ulva* hydrolysate in comparison to that grown on sugar mixture are shown in Figure 8 and 9. The ^1^H-NMR spectral data matched with the ^1^H-NMR spectrum of P(3HBV) acquired by (Bloembergen et al., 1989). From the calculated peak integration, it can be concluded that the PHA produced from *Cobetia* isolate no. 105 grown on sugar mixture (i.e. glucose, fructose and mannitol) contains mainly 3HB with 0.94 mole% 3HV while 3.29 mole% 3HV was obtained when *Cobetia* isolate no. 105 was grown on *Ulva* sp. acid hydrolysate.

**Table 5.** Analysis of sugars obtained by acid hydrolysis of *Ulva* sp.

| Glucose mg/g | Rhamnose mg/g | Galactose mg/g | Xylose mg/g | Fructose mg/g | Glu acid mg/g | Total mg/g |
| --- | --- | --- | --- | --- | --- | --- |
| 16.1±0.8 | 6.2±0.45 | 1.0±0.11 | 1.6±0.22 | 2.8±0.41 | 1.3±0.11 | 27.1±1.83 |

**Figure 7.**
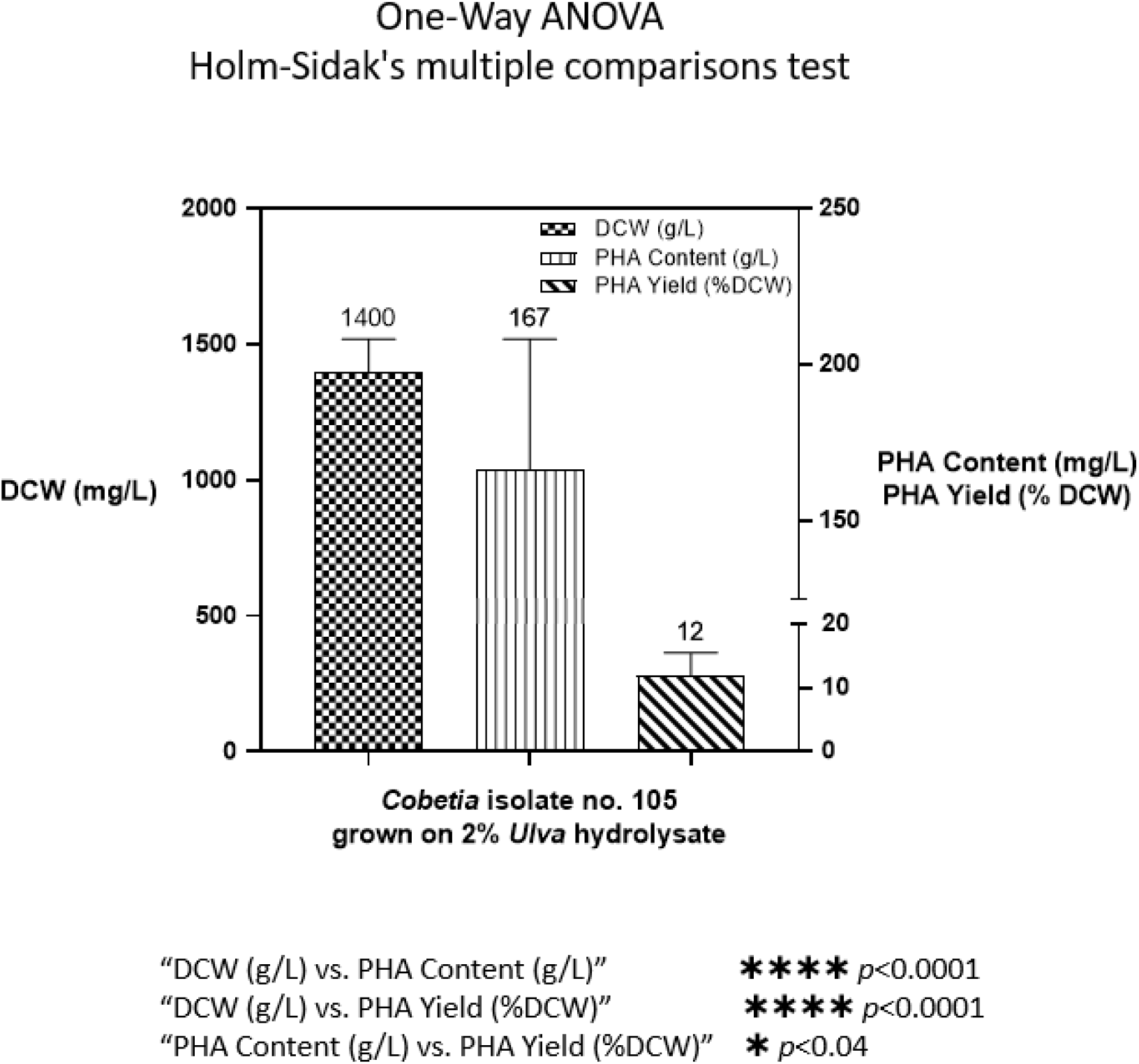
PHA content, PHA yield and DCW of *Cobetia* isolate no. 105 grown on Ulva sp. hydrolysate as a sole carbon. One-Way ANOVA, Holm Sidak’s multiple comparison test was performed.

**Figure 8.**
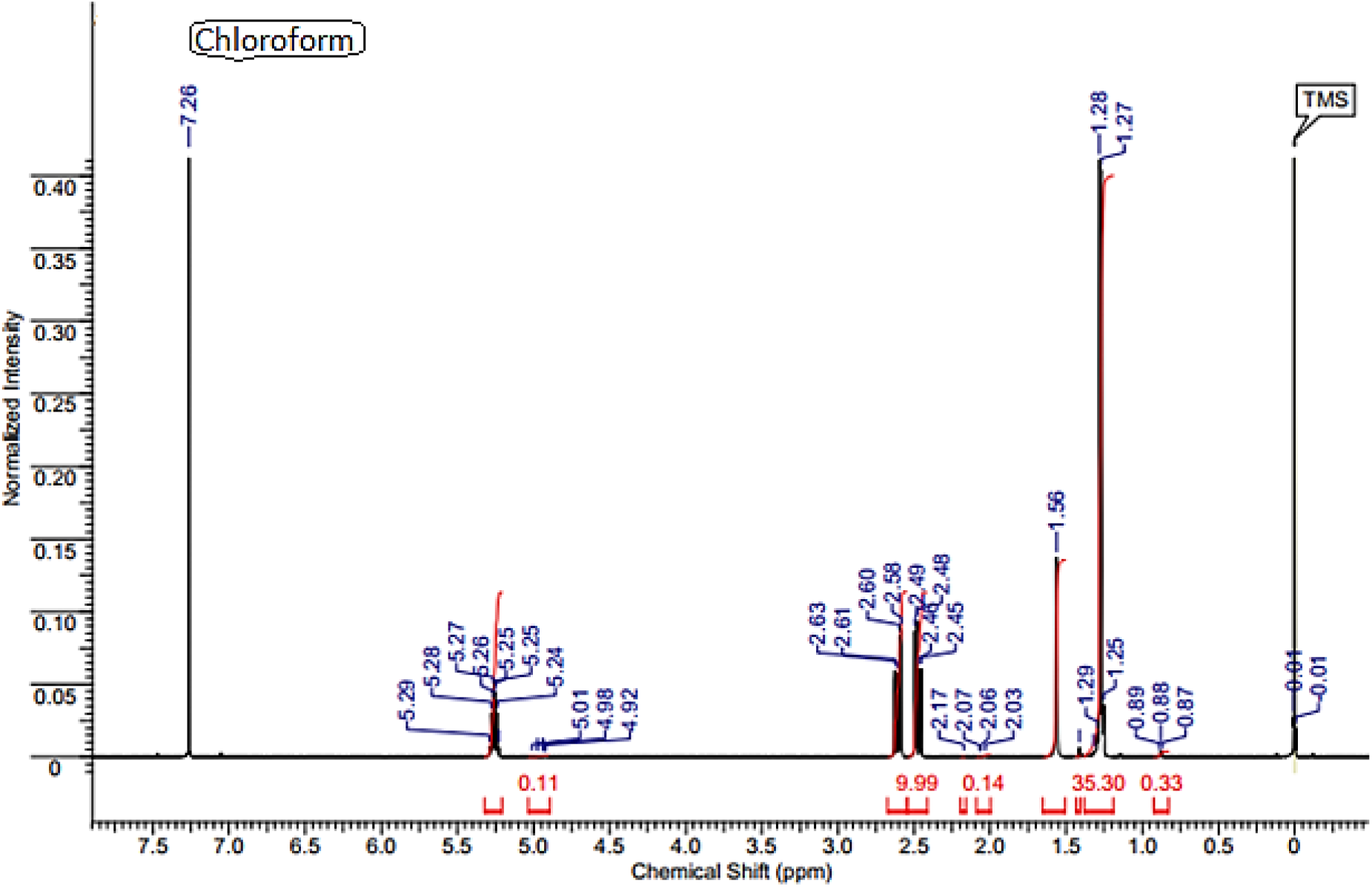
^1^H NMR spectrum of the P(3HBV) produced by *Cobetia* isolate no. 105 when grown on sugar mixture, i.e. glucose, fructose and mannitol and extracted with chloroform.

**Figure 9.**
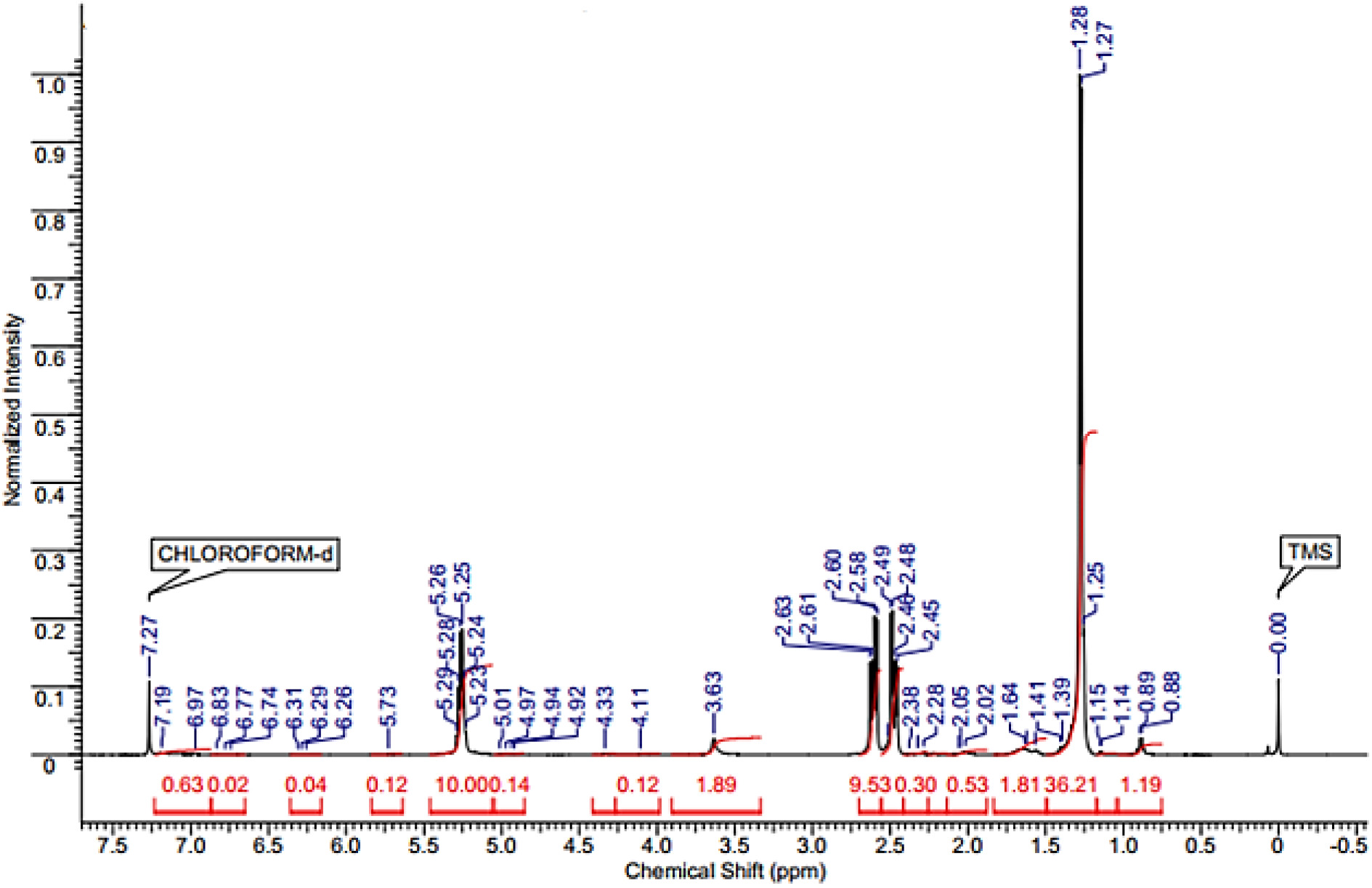
^1^H NMR spectrum of the P)(3HBV) produced by *Cobetia* isolate no. 105 when grown on *Ulva* sp. hydrolysate and extracted with chloroform.

## Discussion

We have successfully isolated bacteria strains from the green seaweed *Ulva sp*. that found to accumulate P(3HB) on various sugars found in seaweed. Taxonomic studies and 16S rDNA gene sequence analysis revealed that these bacteria are phylogenetically related to species of the genus *Cobetia, Bacillus, Sulfitobacter* and *Pseudoaltermonas. Bacillus* isolate no. 3, found to have a close genomic relationship to *B*. cereus, *B. mobilis, B. pacificus* and *B. thuringiensis* with 98% identity. Isolate no. 48 found to have a close relation to *Sulfitobacter* sp. with 98% identity. Isolate no. 71 found to have a close relation with *Pseudoaltermonas* sp. with 98.52% identity. The closest species of *Cobetia* are *C. amphiletci 46-2* with 100% identity, *C. litoralis KMM 3880* with 99.91%, *C. marina* and *C. pacifica* with 99.57%.

The fermentation process was carried out using different bacteria isolates and different sugars for PHA production. Among all the isolates, the highest production of PHA was obtained by *Cobetia* isolate no. 105 on mannitol with 61% of P(3HB), *Cobetia* isolate no. 107 on fructose with 27.5%, *Cobetia* isolate no. 104 on fructose with 23.3%, *Cobetia* isolate no. 75 on glucose with 20.9% and *Cobetia* isolate no. 65 on mannitol with 17.1%.

*Cobetia* strains found to assimilate various carbon sources, such as fructose, glucose, mannitol and galactose, and produce P(3HB) with high productivity. *Cobetia* was classified originally as *Arthrobacter marinus* by (Cobet et al., 1971), then *Deleya marina* by (BAUMANN et al., 1983) and *Halomonas marina* by (Dobson and Franzmann, 1996). The genus *Cobetia* contains mainly two well-known strains, *Cobetia marina* (Arahal et al., 2002), and *Cobetia crustatorum* (Kim et al., 2010).

Several studies were conducted on microbial species genetically related to the *Halomonas* or *Cobetia* genus as PHA-producers such as *Cobetia marina, Halomonas boliviensis LC1, Halomonas elongate DSM 2581, Halomonas salina* and *Halomonas sp*.*TD01* (Mothes et al., 2008; Quillaguamán et al., 2005; Tao et al., 2017).

Among all *Cobetia* strains reported in this study, *Cobetia* no. 105 which was identified as *C. amphiletci* showed the highest P(3BH) yield; 61% w/w when grown on mannitol and 12% w/w on *Ulva* acid hydrolysate as a sole carbon source. Very recently, Moriya et al. (2020) have reported the production of PHB (13.5%) by *Cobetia* strain (5-11-6-3) in a medium containing crushed waste *Laminaria* sp., (brown seaweed). Furthermore, they have used alginate as a substrate for *Cobetia* strain 5-11-6-3 which yielded 62.1% of PHB with a content of 3.11 g L^-1^.

Wang et al. (2010) have evaluated the content of P(3HB) by *Pseudoaltermonas* sp. SM9913 when was grown on glucose, decanoate and olive oil. The strain revealed P(3HB) accumulation of 3.10, 1.89, 2.57% of the cell dry weight when glucose, decanoate and olive oil were provided as a carbon source, respectively (Wang et al., 2010). In our hands, P(3HB) production up to 7.46% (w/w) was obtained by *Pseudoaltermonas* isolate no. 71 on fructose. Mereuta et al. reported for the first time the production of P(3HB) by *Sulfitobacter* genus, which was isolated from the black sea (Mereuta et al., 2018).

The P(3HB) productivity reported for mixed cultures were found to be lower than the productivity of pure cultures (Serafim et al., 2008). The maximum cell concentration reported for aerobic dynamic feeding (ADF) operated systems was 6.1 gL^-1^ (Dionisi et al., 2006), which is much lower than those obtained by pure cultures, usually above 80 gL^-1^ (Lee et al., 1999). The reason for this result is the apparent difficulty in reaching high biomass concentrations in the mixed-culture process (Oehmen et al., 2014), probably due to bacterial competition on the carbon source. Many studies have demonstrated the production of P(3HB) from various marine bacteria (Mostafa et al., 2020a, 2020b; Pu et al., 2020). In this study, some bacteria isolates found to produce P(3HBV).

## Conclusions

In this study, we succeeded in isolating different bacteria strains, including P(3HB) and P(3HBV)-producing bacteria associated with seaweed *Ulva* sp. designated *C. amphiletci, Sulfitobacter* and *Pseudo-altermonas* from various sugars. The selection of a suitable substrate is an important factor for improving microbial PHA production yield, composition and properties. Based on our findings, we recommend conducting large-scale assays and evaluating the industrial production of P(3HB) using these strains in green seaweed biorefineries.

## Authors contributions

A.G. and M.G. conceived the idea of the study. R.G., M.P. and A.G. designed the research experiments. M.P. isolated the bacteria from the seaweed. R.G. performed the experiments and analysed the results. J.S. and M.P. helped with molecular identification. R.U. helped with GC-MS analysis. R.G. and A.G. wrote the manuscript.

## Acknowledgement

R.G. thanks the TRDC-TAU collaborative research grant, the Arianne de Rothchild Women’s Doctoral Program Scholarship, and the Lewis and Martin Whitman Scholarship for Arab students for financial support of this work. The authors thank The Aaron Frenkel Air Pollution Initiative at Tel Aviv University for financial support.

